# RápidoPGS: A rapid polygenic score calculator for summary GWAS data without a test dataset

**DOI:** 10.1101/2020.07.24.220392

**Authors:** Guillermo Reales, Elena Vigorito, Martin Kelemen, Chris Wallace

## Abstract

**Motivation:** Polygenic scores (PGS) aim to genetically predict complex traits at an individual level. PGS are typically trained on genome-wide association summary statistics and require an independent test dataset to tune parameters. More recent methods allow parameters to be tuned on the training data, removing the need for independent test data, but approaches are computationally intensive. Based on fine-mapping principles, we present RápidoPGS, a flexible and fast method to compute PGS requiring summary-level GWAS datasets only, with little computational requirements and no test data required for parameter tuning.

**Results:** We show that RápidoPGS performs slightly less well than two out of three other widely-used PGS methods (LDpred2, PRScs, and SBayesR) for case-control datasets, with median r^2^ difference: −0.0092, −0.0042, and 0.0064, respectively, but up to 17,000-fold faster with reduced computational requirements. RápidoPGS is implemented in R and can work with user-supplied summary statistics or download them from the GWAS catalog.

**Availability and implementation:** Our method is available with a GPL license as an R package from <u>GitHub</u>.

## 1. Introduction

Genome-wide association studies (GWAS) have been widely successful at identifying a large number of genetic variants (usually single nucleotide polymorphisms, or SNPs) associated with a wide range of diseases and complex traits (Buniello *et al.*, 2019). Most genetic variants have a small individual effect on the tested traits and thus have low predictive power (Yang *et al.*, 2010; Dudbridge, 2013; International Schizophrenia Consortium *et al.*, 2009). However, simultaneously evaluating the effects of all common SNPs has the potential to explain much of the heritability for complex diseases and phenotypes (Yang *et al.*, 2010; Bulik-Sullivan *et al.*, 2015).

Polygenic scores (PGS) estimate individual propensity to a phenotype (e.g. disease) by summing an individual’s genotypes (coded 0, 1, 2), weighted by their effect sizes, estimated from GWAS data

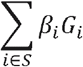

where *β_i_* is the estimated effect for variant *i*, and *G_i_* the genotype. Challenges relate to the nature of GWAS data to determine the set of SNPs *S* to use. If ignored, linkage disequilibrium (LD) between variants would mean double-counting the effects of causal variants in high LD with multiple other variants. Additionally, error in the estimated effect sizes is most pronounced for small effects, for which true association may not be distinguishable from noise about a null value. Different approaches have been developed to deal with these. Initial approaches selected the most strongly associated variant in each genome wide-significant peak. However, it was soon realised that predictive accuracy could be improved by reducing the significance threshold, to capture more truly associated variants even at the cost of including some false associations (which should add noise, but not bias, to any prediction), especially for highly polygenic diseases (Chatterjee *et al.*, 2016; Dudbridge, 2013). Methods were developed to use an external test set to tune the significance threshold parameter, as well as to use automated LD-pruning algorithms to select independent variants rather than selecting one variant per peak (Privé *et al.*, 2019; Euesden *et al.*, 2015).

More statistically sophisticated approaches have since been developed to average over multiple variants in LD without double counting (which is more accurate than selecting just one), shrinking effect estimates by a continuous weight, *w_i_* (Privé, Arbel, and Bjarni J Vilhjálmsson, 2020; Vilhjálmsson *et al.*, 2015; Ge *et al.*, 2019)

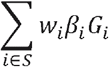

As with the “prune and threshold” approach, such methods initially required external independent test datasets to tune their parameters, which can present a barrier to practical use. Recently, automated methods have been developed that remove the need for an external test dataset via hierarchical Bayesian models (Privé, Arbel, and Bjarni J Vilhjálmsson, 2020; Ge *et al.*, 2019), and perform nearly as well as their externally tuned counterparts. However, they require storage and inversion of large LD matrices, which is computationally intensive, and tuning, internally or externally, adds a further layer of iterative computation. Some approaches mitigate this by adding a thinning step, discarding a subset of SNPs to reduce the burden (e.g. LDpred2 recommends restricting SNPs to those in the HapMap3 panel, Privé, Arbel, and Bjarni J Vilhjálmsson, 2020), but constructing PGS still takes a long time.

Our framework is based on considering PGS construction as a fine-mapping problem. If we knew exactly the causal variants for a trait, an optimal PGS would comprise the estimated effects of just those variants. Fine-mapping methods estimate probabilities that a variant is causal given observed data. So, in the absence of knowing the exact set of true causal variants, a natural PGS might be constructed as above, with *w_i_* set to the probability variant *i* is causal - i.e. by replacing the estimated effect size of each SNP by the posterior expectation of its causal effect. Whilst modern PGS methods focus on the accurate estimation of SNP effects in a joint model, the optimal solution to the joint model is also that which puts non-zero effects only on the true causal variants. We note that many of these approaches involve estimating the probability that a specific SNP has a non-zero effect - the estimand itself in fine-mapping - but the overarching goal remains the estimation of SNP effects. Our intention in focusing directly on the probabilities of causality is to allow us to explore whether established fine-mapping tools can be adapted to the PGS question, benefiting from their speed relative to existing PGS methods.

We present two approaches that can accommodate different assumptions on the number of causal variants in each LD-defined region. The basic fine-mapping approach is very fast, because it makes a simplifying assumption that only a single causal variant may exist in any LD-defined region, based on the principles developed by Maller *et al.*, (2012). While unrealistic, it has been shown to perform well for fine-mapping and, of relevance to PGS, allows the definition of posterior probabilities of causality using predefined LD regions and without the burden of processing large LD matrices (Maller *et al.*, 2012). Distinguishing noise around null associations from true associations is dealt with by two parameters: a prior probability that a random SNP is causal and the variance of the prior distribution of effect sizes at true causal variants. While in a PGS these would be open to tuning, in fine-mapping sensible default values can be chosen that reflect existing knowledge from the breadth of GWAS studies already conducted (Wallace, 2020).

It is possible to relax the single causal variant assumption if additional information on LD is available. We propose using an alternative fine-mapping method based on the “sum of single effects” model (SuSiE, Wang *et al.*, 2020). SuSiE is a computationally efficient approach to variable selection in linear regression, which uses a multiple causal variant model to compute the posterior inclusion probability for each variant in an LD-defined region, which we can use as *w_i_*.

We present RápidoPGS, a lightweight and fast (*rápido*, in Spanish) method to compute PGS based on fine mapping approaches that only requires a summary statistics dataset, with no need for an independent test dataset to adjust parameters, and that can generate weights for millions of SNPs in only a few seconds or minutes. RápidoPGS works fully in R, requiring few dependencies. A PGS can be quickly constructed from any GWAS summary statistic dataset, or any GWAS PubMed ID, if harmonised datasets are available at GWAS catalog. We created PGS models for eight case-control and two quantitative traits using RápidoPGS and three widely used PGS methods for comparison: LDpred2 (Privé, Arbel, and Bjarni J Vilhjálmsson, 2020), PRScs (Ge *et al.*, 2019), and SBayesR (Lloyd-Jones *et al.*, 2019). We then used UK Biobank data as a validation dataset to evaluate the predictive performance of all models, and benchmarked their run times.

## 2. Materials and Methods

### 2.1. Overview of our approach

We compute posterior probabilities of causality at each variant within LD-blocks defined by lddetect (Berisa and Pickrell, 2016) (publicly available at https://bitbucket.org/nygcresearch/ldetect-data/src).

Under a single causal variant assumption, given summary statistics at each SNP *i*, *β_i_*, *V_i_* (respectively, the estimated effect size and its variance), we can define an approximate Bayes factor summarising the evidence for its association (Wakefield, 2009) as

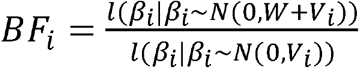

where l is the likelihood of seeing the observed value of *β_i_* conditional on it following a normal distribution with specified variance, *N* is a normal distribution, and *W* is the variance of the prior on the true effect size. We assume that the true causal effect size at any SNP is sampled from an *N*(0,*W*) distribution.

Under the at most one causal variant assumption (often called a single causal variant assumption, but it includes the possibility of no causal variant), the posterior probability that SNP *i* is causal in a region of *n* SNPs, is

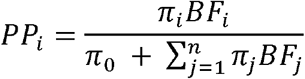

where *π_i_* is the prior SNP *i* is causal (typically set to the same value for all SNPs) and *π*_0_ =1 − *nπ_i_* is the prior probability there is no causal variant in the region (Maller *et al.*, 2012). Note therefore 0 ≤ *PP_i_* < 1 and Σ_*i*_ *PP_i_* ≤ 1, so that *PP_i_* can serve to shrink SNP effect estimates more or less according to the posterior belief that a SNP is causal for a trait.

We set default parameter values for RápidoPGS according to currently widely adopted values: *π_i_* = 10^−4^, *W* = 0.2^2^ for case-control traits (Wallace, 2020). There is less consensus on appropriate values of *W* for quantitative traits, and we propose to estimate *W* given estimated heritability of the trait under study or a similar trait, which is often available (e.g. at LDhub, http://ldsc.broadinstitute.org/ldhub/, Zheng *et al.*, 2017):

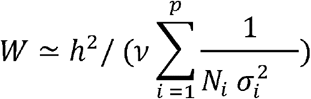

where *h*^2^ is the estimated trait heritability, *v*is the prior that a given variant is causal (we set *v*= 10^−4^), *p* is the number of variants, *N_i_* is the number of individuals used in the inference for variant *i*, and 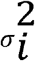 is the variance of the estimated effect *β_i_* associated with variant *i in the GWAS summary statistics for the trait of choice*. See Supplementary Note for the full derivation of the equation.

Alternatively, we can relax the single causal variant assumption. SuSiE (Wang *et al.*, 2020) is an approach for variable selection in regression, based on previous models for Bayesian variable selection in regression (BVSR), but with a different structure that allows for a faster and simpler model fitting procedure based on fitting multiple ‘single-effect regression’ models (multiple-regression models with exactly one variable with non-zero regression coefficient), and then constructing the overall effect vector as the sum of the single-effects vectors (hence, sum of single effects model). SuSiE requires the same parameters (*π_i_* and *W*) as the single causal variant approach, although *W* may be internally estimated. We call the two approaches to estimating *W* “informed” or “auto”.

Finally, we considered pruning input SNPs according to their p values using 2 parameters in order to increase speed: (1) α_block_, the minimum p-value at least one variant in a LD block must have for the block to be considered for subsequent analyses (i.e. if no SNPs in a block have p-values below α_block_, the entire block is skipped), and (2) α_SNP_, the minimum p-value a variant within a selected block must have to be considered.

### 2.2. GWAS datasets and SNP QC

We downloaded ten publicly available GWAS summary statistics datasets on the following traits: eight case-control datasets: asthma (Demenais *et al.*, 2018), rheumatoid arthritis [RA] (Okada *et al.*, 2014), type 1 diabetes [T1D] (Cooper *et al.*, 2017), type 2 diabetes [T2D] (Scott *et al.*, 2017), breast cancer [BRCA] (Michailidou *et al.*, 2017), prostate cancer [PRCA] (Schumacher *et al.*, 2018), coronary artery disease [CAD] (Nikpay *et al.*, 2015), and major depression disorder [MDD] (Wray *et al.*, 2018). We also added two quantitative traits: body mass index [BMI] (Locke *et al.*, 2015) and height (Wood *et al.*, 2014). All datasets considered are meta-analyses of GWAS performed on European populations, with the exception of RA, which is a trans-ethnic meta-analysis including European and Asian ancestries.

We applied common quality control to all datasets prior to PGS generation by removing unneeded rows with missing data at required columns (genomic coordinates, alleles, allele frequencies, log(OR), standard error), and from case/control sample size column, if provided. Following LDpred2 quality control guidance for summary statistic datasets, we computed the effective sample size (N_eff_ = 4 / (1 / N controls + 1 / N cases)), the summary statistic standard deviation (SD_ss_), defined as 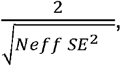, , where SE is the standard error of the effect for each SNP, and the validation standard deviation (SDval), which here we computed as 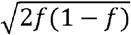, where *f* is the allele frequency for the effect allele. When this frequency was not available in the dataset, we computed it using the CEU population in 1000 Genomes Phase III panel. Note that in the LDpred2 paper, SDval is computed using the test dataset, but since our aim is not to use any external dataset for QC or parameter tuning, we computed it as above. We then excluded SNPs for which SDss < 0.5 ◻ SDval or SDss > SDval + 0.1 or SDss < 0.1 or SDval < 0.05, following LDpred2 guidelines (Privé, Arbel, and Bjarni J Vilhjálmsson, 2020). We plotted SDss vs. SDval and visually inspected the relationship to spot possible deviations (Supplementary Fig. 1–10).

We further filtered SNPs to the subset overlapping the HapMap3 variants, as recommended by LDpred2, PRScs, and SBayesR, the three methods we compared RápidoPGS to. To explore how RápidoPGS prediction ability changes depending on the selection of SNPs, we created another set of post-QC datasets, filtered by 1000 Genomes Phase III variants instead.

### 2.3. PGS generation using RápidoPGS

RápidoPGS comes in two flavours: RápidoPGS-single, and RápidoPGS-multi. As explained in section 2.1, they correspond to two different approaches, which differ in their assumption of how many causal variants per LD block are considered in their respective models — hence “single” and “multi”. RápidoPGS-single requires specifying *W*, which for case-control traits we set as W = 0.2^2^. For quantitative traits, we estimated the informed *W* using estimates of heritability from LDhub: “BMI” *h*^2^ = 0.246, and “Standing Height” *h*^2^ = 0.462 which led to estimates of *W* of (0.127)^2^ for BMI, and (0.141)^2^ for height. RápidoPGS-multi allows for W to be automatically estimated, and it is our recommended choice.

All methods included in this study (with the exception of RápidoPGS-single) require LD matrices. LD matrices inform the method about the correlations between SNPs in a reference population, which is ideally close to the population we create the PGS for. LDpred2, SBayesR and PRScs offer European HapMap3-filtered LD matrices, which are supplied with each method. RápidoPGS-multi can either use a reference panel to compute LD matrices or input pre-computed LD matrices supplied by the user. In this study, we employed UKBB-based LD matrices provided by LDpred2 authors, publicly avaliable at https://figshare.com/articles/dataset/European_LD_reference/13034123.

To improve speed and reduce computational cost for RápidoPGS-multi, we thinned the input SNPs using α_SNP_ = 0.1, or α_SNP_ = 0.01. As only 10% of truly null SNPs achieve p values < 0.1 and only 1% < 0.01, this should remove 90% or 99% of null SNPs respectively. Assuming most blocks have 1000 SNPs or more, nearly all blocks containing only null SNPs should achieve a minimum p value of 10^−3^, and only 10% a minimum p value of 10^−4^, so we chose block-thinning parameters of α_block_ = 10^−3^ or α_block_ = 10^−4^. Of note, although we used 15 CPUs and 8 hours as a standard across all methods, RápidoPGS-single and RápidoPGS-multi (using pre-computed LD matrices) do not require using multiple CPUs, and our tests showed that one CPU is sufficient to run the method, without speed lost compared with using multiple CPUs. This makes RápidoPGS especially suitable for computational environments with limited resources.

### 2.4. PGS generation using LDpred2-auto

For PGS model generation using LDpred2-auto (Privé, Arbel, and Bjarni J Vilhjálmsson, 2020) we followed the LDpred2 instructions and adapted the code provided in the tutorials and the accompanying code (https://github.com/privefl/paper-ldpred2 and https://privefl.github.io/bigsnpr/articles/LDpred2.html). However, we omitted the prediction steps, as our approach does not consider a test dataset for PGS model generation (see companion code to this paper for implementation details).

We ran LDpred2-auto on a per-chromosome basis for all 22 autosomes. LDpred2 uses two essential hyperparameters to compute the adjusted effect sizes: p (the proportion of causal variants), and h2 (the heritability of the trait). LDpred2-auto can estimate both hyperparameters from the training data, thus not requiring a test data set to tune them. For p, it can take multiple a number of initial values, from which Gibbs sampling chains run for a fixed number of iterations to find the optimal effect sizes for each SNP. After 4,000a number of iterations (3000 after 1000 burn-in), we averaged betas across all chains. As recommended by LDpred2 authors in the LDpred2 tutorial (https://privefl.github.io/bigsnpr/articles/LDpred2.html), we used 15 initial values for p, ranging from 10^−4^ to 0.9. We computed initial h2 from the data using the snp_ldsc function. To ensure convergence, we used 4,000 iterations (3,000 iterations + 1,000 burn-in). We skipped computation for chromosomes for which estimated h2 was below 1e-4, as was done in the LDpred2 paper (Privé, Arbel, and Bjarni J Vilhjálmsson, 2020). Note that LDpred2 authors recommend running LDpred2 genome-wide rather than per-chromosome, due to better performance in their tests. However, LDpred2-auto genome-wide approach did not finish on time for all but one trait (T1D), using 32 hours and 15 CPUs on our HPC. The two approaches for that trait gave very similar results (difference in r^2^ = 0.0031). Our LDpred2 per-chromosome are also very similar to the reported genome-wide results in LDpred2 paper (P^r^ivé, Arbel, and Bjarni J Vilhjálmsson, 2020). Therefore, we report results for the per-chromosome approach.

### 2.5. PGS generation using PRScs-auto

We ran PRScs (Ge et al., 2019) using pre-computed LD matrices from European 1000 Genomes Project phase 3 samples, as provided by the authors. We formatted the input summary statistic datasets following documentation and ran PRScs-auto using default parameters. PRScs requires GWAS sample size (N), so we used case + control numbers for case-control datasets. MDD dataset had per-SNP N, so we used the sum of median cases and median controls (which were also max values). For BMI and height datasets, which also had per-column N, we used the median value of each dataset as its N (233,691 and 252,021 respectively). We ran PRScs for all chromosomes, and concatenated individual files together. Although it was unclear if PRScs would benefit from parallel computation, we ran it with 15 CPUs and 8 hours for consistency.

### 2.6. PGS generation using SBayesR

SBayesR is part of the GCTB suite (Lloyd-Jones *et al.*, 2019). SBayesR requires LD matrices from a reference panel too, so we downloaded shrunk sparse LD matrices described in Lloyd-Jones et al. (2019), computed based on a random sample of 50K individuals of European ancestry in UK Biobank data and a genetic map in the public domain, following the algorithm in Wen and Stephens (2010). These matrices comprise the HapMap3 variants, like our filtered datasets. SBayesR requires effect allele frequency, and we precomputed them using the CEU population in a 1000 Genomes Phase III panel. We used the same strategy for supplying sample size as we did for PRScs. We experienced some issues related to poor convergence, so following SBayesR documentation advice, we dropped SNPs in the lower 10% N quantile for the three datasets with per-SNP N. We also used the --imputeN flag to allow GCTB estimate per-SNP sample size based on the beta values, SE and allele frequencies and exclude SNPs that have the imputed sample sizes 3 standard deviations apart from the median value. In addition, we used additional --p-value 0.5 --rsq 0.99 flags to remove SNPs with P-values above 0.5 and with r^2 larger than 0.99, so mitigating the effect of SNPs in high LD with opposite effects. We ran SBayesR with default values, 15 CPUs and 8 hours. We ran SBayesR for all chromosomes in one go, using --mldm tag and a list of matrices for each chromosome. All datasets ran successfully except for T1D, which despite the measures mentioned above continued to experience convergence issues and failed to provide an output.

### 2.7. Model evaluation using UK Biobank individual data as a validation dataset

We evaluated the predictive performance of our PGS models generated with the four methods using individual genotypes from the UK Biobank cohort. For all traits we excluded SNPs with low imputation quality (info score <0.3) or multi-allelic SNPs. Moreover, we removed related individuals and restricted the analysis to individuals of European ancestry (UKBB field 22006, genetic ethnic group). The binary traits selected (Table S1) were the same used in the evaluation of LDpred2 (Privé, Arbel, and Bjarni J. Vilhjálmsson, 2020) and we applied the same selection criteria for cases and controls as previously described (Privé *et al.*, 2019) following the code for each of the traits as in https://github.com/privefl/simus-PRS/tree/master/paper3-SCT/code_real). For all disease traits we included as cases those individuals who self-reported the condition or were diagnosed by a medical doctor or the condition was included in their death record. For breast and prostate cancer we excluded individuals with other cancer diagnosis. Moreover, for breast cancer we restricted the analysis to females, for prostate cancer to males. For rheumatoid arthritis we excluded individuals with any other musculoskeletal system and connective tissue condition. For type 1 diabetes we excluded individuals with type 2 diabetes and vice versa. For coronary artery disease we excluded individuals with other heart conditions. For asthma we excluded individuals with additional respiratory conditions. For MDD we excluded individuals with additional mental and behavioral disorders as well as individuals which were included in the GWAS used to construct the score. The MDD GWAS we used to generate the score contained ~ 30,000 individuals from UKBB which corresponded to the initial release and were genotyped with the BiLEVE array. We identified those individuals using code “22000” and excluded those genotyped with the BiLEVE array (batches coded −1 to −11). For BMI and height we used the UK Biobank codes “21001” and “50”, respectively.

After computing the scores, for case control phenotypes we estimated the area under the curve (AUC) adjusting for the first 40 genetic principal components (PCs, codes “22009-0.1-40”), age (code “21003-0.0”) and sex (code “22001-0.0”) -the latter for all traits except breast and prostate cancer- using the R package “ROCnReg”. We then obtained *r*^2^ on the liability scale using the estimated AUC (Lee *et al.*, 2012) as:

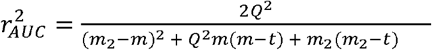

with:

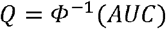

m = mean of liability for cases

K = population prevalence

t = threshold on the normal distribution which truncates the proportion of disease prevalence

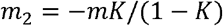

We estimated K as the proportion of cases from the UKBB dataset.

For continuous traits we assessed the squared correlation between the PGS and the measured trait (r^2^) adjusting or the first 40 genetic PCs (codes “22009-0.1-40”), age (code “21003-0.0”) and sex (code “22001-0.0”). Briefly, we regressed each trait against the covariates (PCs, age and sex) and then correlated the residuals with the predictive score for the relevant trait. We constructed 95% confidence intervals for our estimates by bootstrapping 1000 times.

### 2.8. Runtimes

We timed all methods and approaches for all datasets in independent runs, using the same HPC parameters (ie. 15 CPUs, 8 hours). We used system.time() function in R for timing RápidoPGS, LDpred2, and SBayesR wall clock runtime. For PRScs, we used the unix “time” programme, as using system.time() was unfeasible.

Times for RápidoPGS includes full PGS generation procedure: check dataset integrity (for both single and multi), handling of pre-computed LD matrix or LD matrix computing from panel and α filtering (RápidoPGS-multi only), algorithm running, and final weight computation. SBayesR and PRScs require transforming the input data into a specific format, a step we did not include in the timing.

## 3. Results

We computed PGS for 10 different traits (Table 1, Supplementary Table 1). We first assessed the relative performance of each RápidoPGS approach. For RápidoPGS-multi, we considered two pairs of α parameters for all traits. The milder thinning (α_SNP_= 0.1, α_block_= 10^−3^), which discards fewer SNPs, showed slightly better performance than α_SNP_= 0.01, α_block_= 10^−4^ setting. RápidoPGS-multi, which allows for multiple causal variants, achieved better performance than RápidoPGS-single, which is simpler and faster, but has the limitation of assuming a single causal variant per block, which is unrealistic in most scenarios. However, for certain traits (eg. RA and T1D), differences in AUC and r^2^ for both approaches are small (Fig 1). Although W (prior variance on the true effect of a variant) can be provided by the user for RápidoPGS-multi, we found that letting RápidoPGS-multi compute W automatically provides better performance, at the expense of increased running time (Supplementary Tables 2 and 3).

**Table 1.** Training datasets used for PGS computation using four different methods in this study. Validation set refers to the UK Biobank individual data used for PGS evaluation.

| Trait | First author | Year | PMID/doi | Controls training set | Cases training set | Total training sample | Controls validation set | Cases validation set | Total validation sample |
| --- | --- | --- | --- | --- | --- | --- | --- | --- | --- |
| Asthma | Demenais | 2018 | 29273806 | 107715 | 19954 | 127669 | 261974 | 43785 | 305759 |
| RA | Okada | 2014 | 2439034 | 61565 | 19234 | 80799 | 226320 | 5614 | 231934 |
| T1D | Cooper | 2017 | doi:10.1101/120022 | 8828 | 5913 | 14741 | 314535 | 771 | 315306 |
| T2D | Scott | 2017 | 28566273 | 132532 | 26676 | 159208 | 314535 | 14175 | 328710 |
| BRCA | Michailidou | 2017 | 29059683 | 119078 | 137045 | 256123 | 158385 | 11578 | 169963 |
| PRCA | Schoemacher | 2018 | 29892016 | 61106 | 79148 | 140254 | 141551 | 6382 | 147933 |
| CAD | Nikpay | 2015 | 26343387 | 123504 | 60801 | 184305 | 225917 | 12263 | 225917 |
| MDD | Wray | 2018 | 29700475 | 113154 | 59851 | 173005 | 255306 | 22287 | 277593 |
| BMI | Locke | 2015 | 25673413 | - | - | 339224 | - | - | 334527 |
| Height | Wood | 2014 | 25282103 | - | - | 253288 | - | - | 334891 |

**Figure 1.**
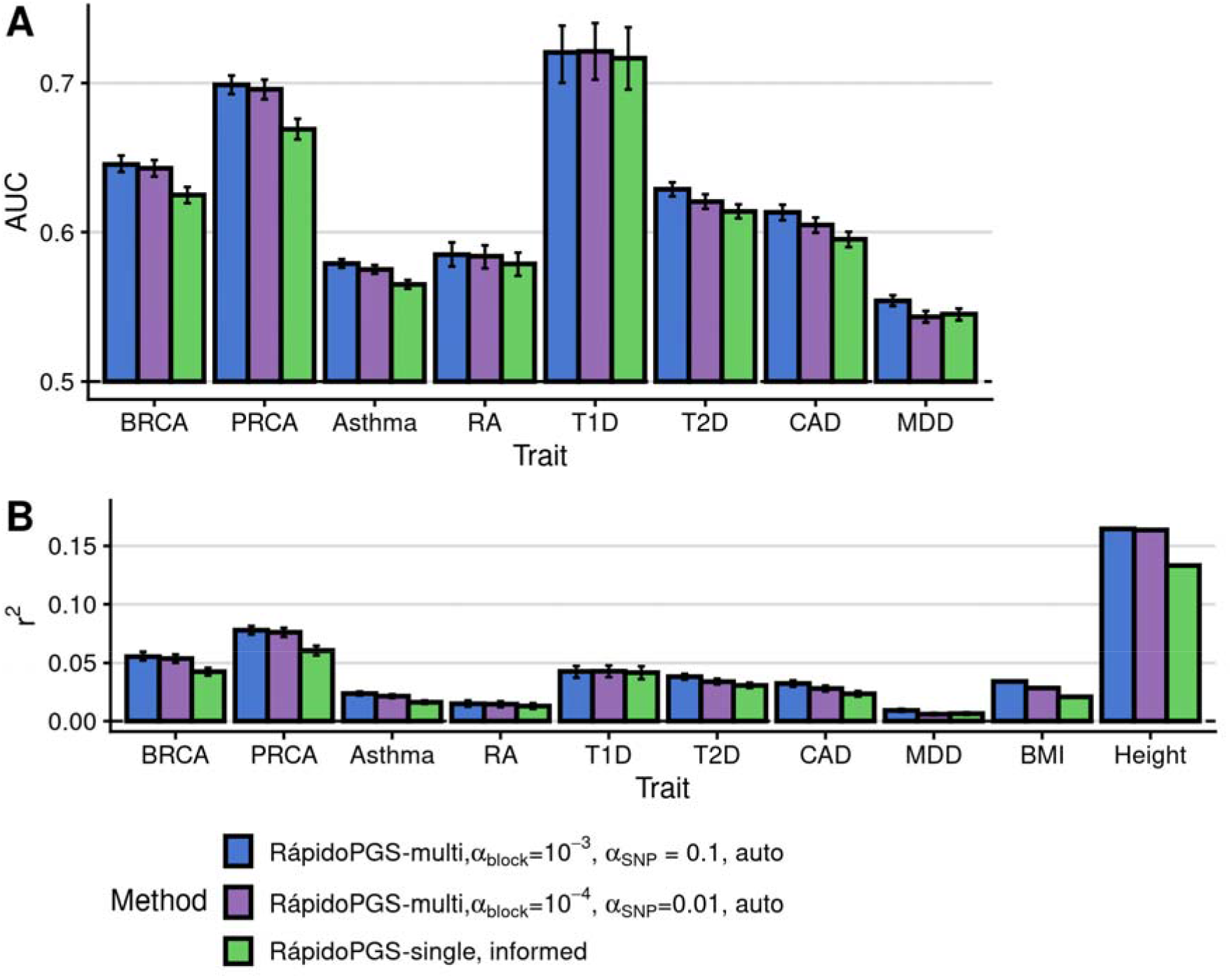
RápidoPGS-multi shows better performance than RápidoPGS-single, with best results when discarding fewer SNPs (α_SNP_=0.1, α_block_=10^−3^), although differences across α parameters are small. A. AUC for case-control traits. B. r^2^ for all traits. In (A) and (B) the error bars correspond to the 95% confidence interval.

We compared the speed of RápidoPGS-single and -multi applying different thresholds, which control the number of input LD blocks and SNPs to construct the PGS (see methods). Selecting α_block_=10^−4^ and α_SNP_=0.01 thresholds which reduces the number of LD blocks and SNPs relative to α_block_=10^−3^ and α_SNP_=0.1, lowered the run time from ~36% (height) to ~64% (asthma) (Fig. 2).

**Figure 2.**
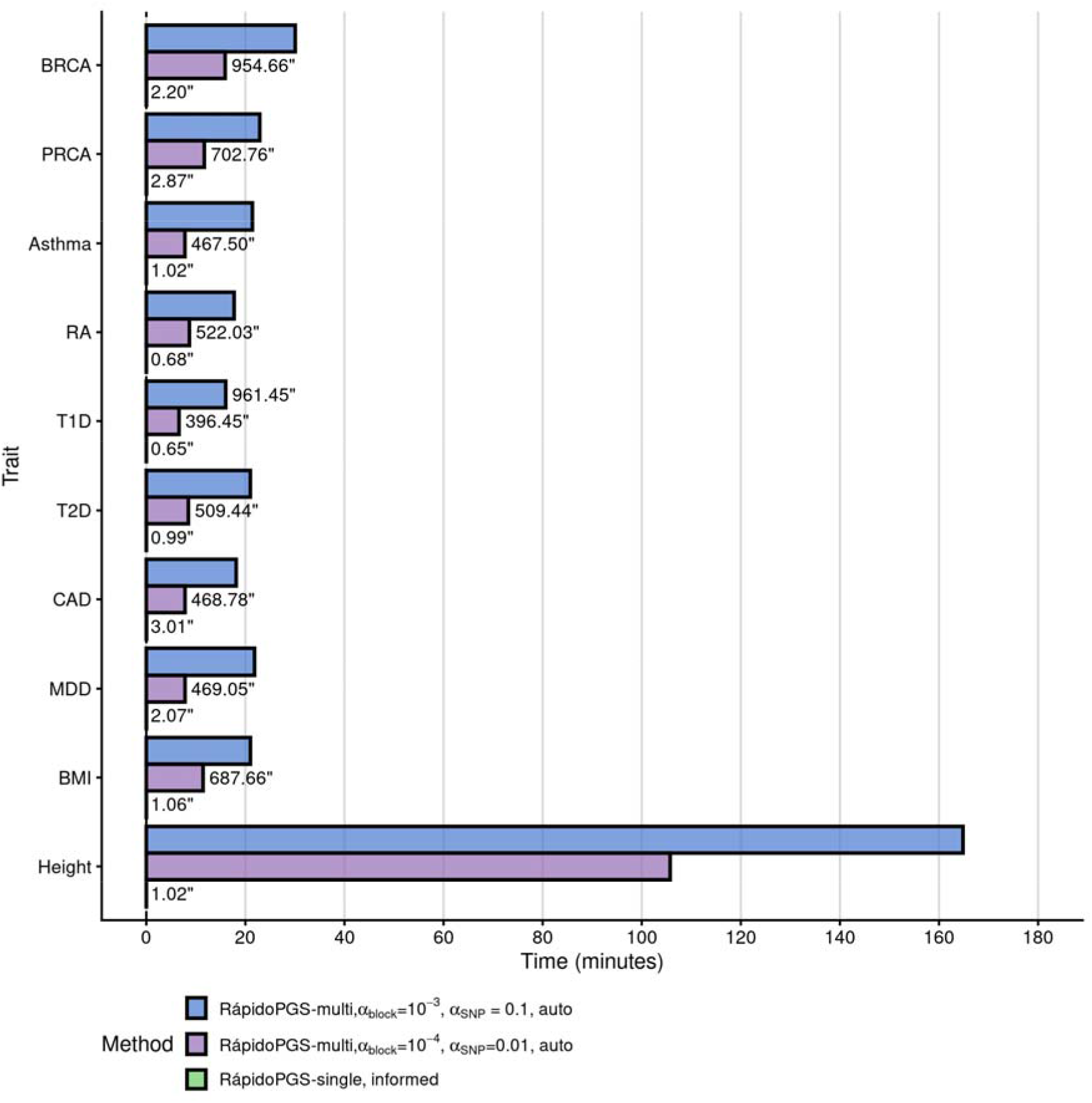
Wallclock times for RápidoPGS approaches (using pre-computed UK Biobank LD matrices and HapMap3 variants), in minutes. Time for runs that took <1,000 seconds is displayed on the right of the bar, in seconds.

We next compared RápidoPGS performance with those of LDpred2, PRScs, and SBayesR. RápidoPGS achieved reasonably good prediction performance for most case control traits, being superior to SBayesR in most instances, and reaching prediction values close to PRScs although lower than LDpred2, which showed the best performance for most traits (Fig 3, Supplementary Table 2). SBayesR experienced convergence issues for T1D that we could not fix. RápidoPGS showed poor performance for MDD and both quantitative traits (BMI and Height).

**Figure 3.**
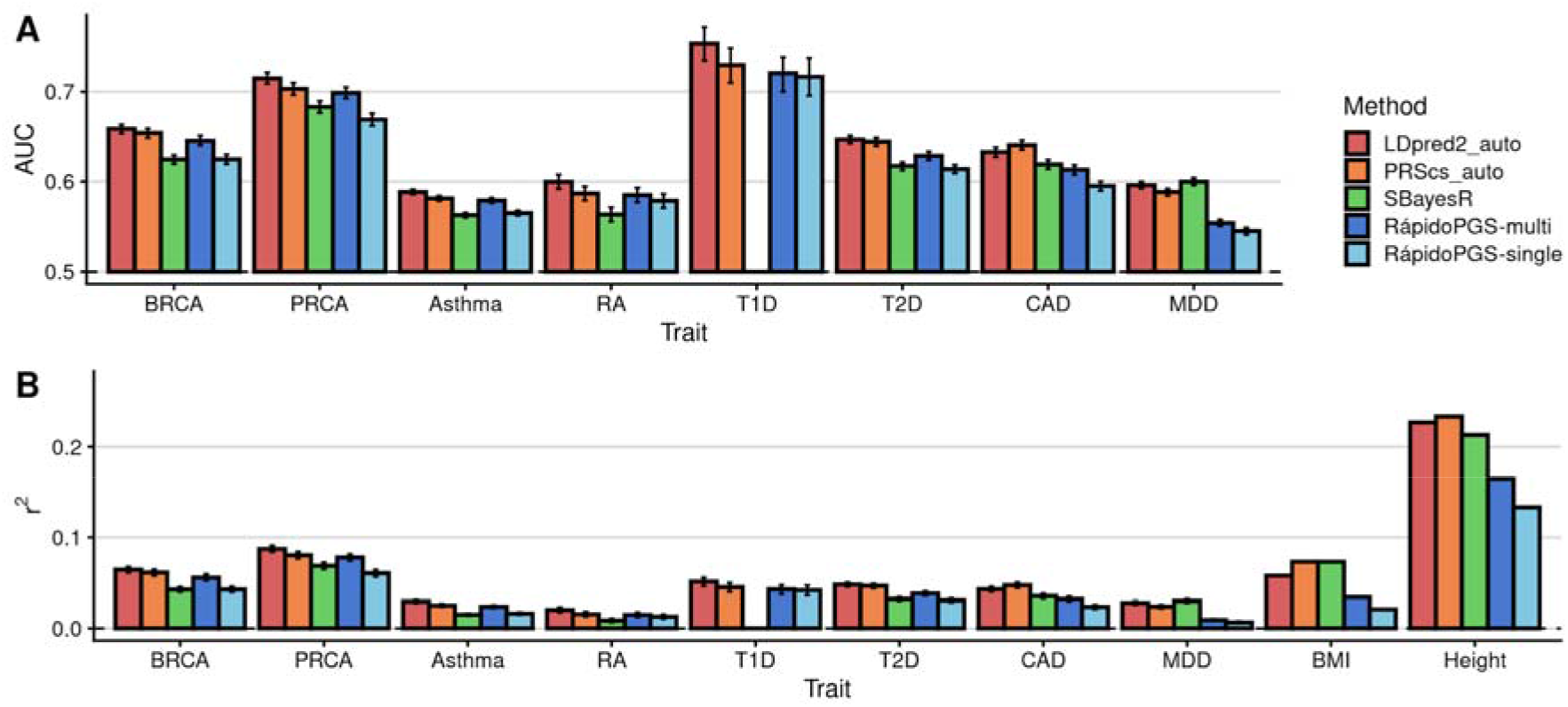
Comparison of RápidoPGS to other methods. A. AUC for case-control traits using LDpred2-auto (per chromosome), PRScs-auto, SBayesR,-auto and RápidoPGS (single and multi with α_block_ = 10^−3^, α_SNP_ = 0.1, and automatic SD prior parameters). For comparison purposes, we added AUC for LDpred2-auto genomewide, reported by (Privé, Arbel, and Bjarni J Vilhjálmsson, 2020). B. r^2^ results for all methods and traits BMI and height. No LDpred2 paper results were available for these traits. SBayesR failed to run for T1D and hence we show an empty column. In (A) and (B) the error bars correspond to the 95% confidence interval.

RápidoPGS-multi (α_SNP_=0.1, α_block_=10^−3^) is the best-performing RápidoPGS method, which shows the smallest median r^2^ difference and median r^2^ ratio to LDpred2 (−0.0092 median difference for case-control, and −0.0429 for quantitative traits, and 0.8011 and 0.6585 median ratio, respectively) and PRScs (mean r^2^ difference = −0.0042, and −0.0538, median r^2^ ratio = 0.9249 and 0.5863), and outperforms SBayesR for case-control traits (mean r^2^ difference = 0.0064 and −0.0436, median r^2^ ratio = 1.1787 and 0.6199) (Supplementary Table 4 and 5).

With regards to runtimes, RápidoPGS-single is the fastest method, being ~8,000-27,000 times faster than the slowest method for each trait (Fig. 4). RápidoPGS-multi approach involves multiple steps, including SNP filtering, LD matrix computation from panel (if not pre-computed), W automatic estimation, and SuSiE algorithm running. All these steps make it inevitably slower than simpler RápidoPGS-single. Nonetheless, RápidoPGS-multi is generally much faster than any of the other methods (1.12-22.17 times faster). We only observed one instance that RápidoPGS-multi took unusually long to finish (Height in Fig. 2 and 4), due to the SuSiE internal algorithm needing to run many iterations to converge in some LD blocks.

**Figure 4.**
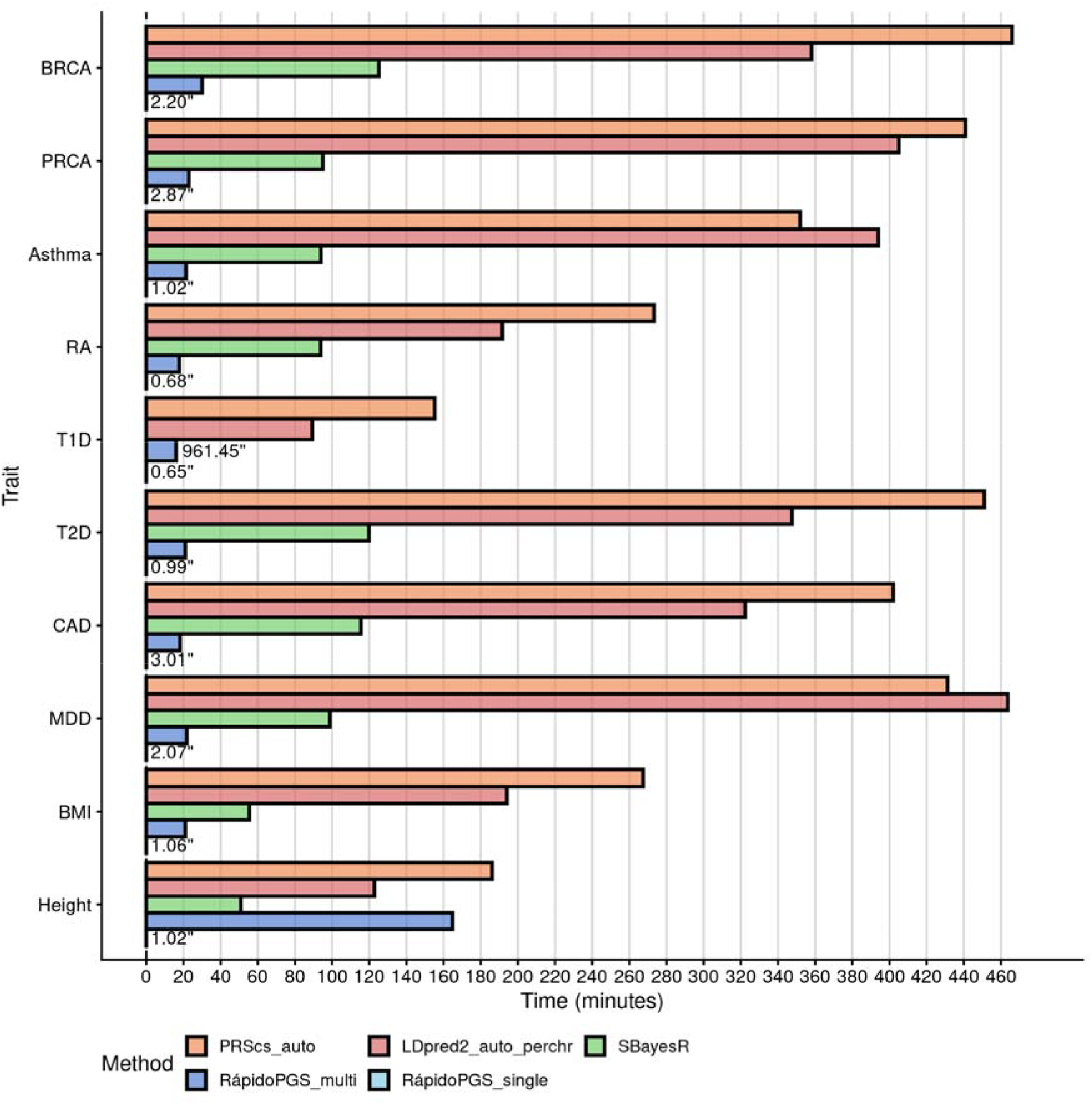
Wallclock times for all methods applied in this study, in minutes. As in Figure 1, RápidoPGS-multi run is represented by α_block_ = 10^−3^, α_SNP_ = 0.1, and automatic SD prior parameters. For better visualisation, running times for runs that took <1,000 seconds are displayed on the right of the corresponding bar, in seconds.

## 4. Discussion

The main downside of most sophisticated PGS methods is their computational cost and running time, taking many hours to finish even when using multiple cores in a high performance computing context.

Having PGS scores computed easily and quickly can be advantageous in a context in which there is a need for rapid assessment of many traits. For example, PGS can be used to estimate “genetic nurture” effects on trait values (Balbona *et al.*, 2021) and a search for traits affected by genetic nurture might be more efficient using a two stage approach: rapid assessment using a RápidoPGS approach, followed by a more accurate estimate of that genetic nurture for selected traits using a more accurate PGS, such as LDpred2.

Alternatively, in simulation studies to explore the utility of PGS for novel applications, a fast but less accurate method may be preferred for practical reasons.

We have thus developed a method to fill this gap, which comprises two flavours, based on fine-mapping strategies with different assumptions. On the basis of the traits considered here, we recommend using RápidoPGS-multi, as it performs generally better than RápidoPGS-single. For RápidoPGS-multi, we recommend allowing internal estimation of W, as it showed to outperform user-supplied W in most cases (Supplementary Table 1). However, we recommend that when suitable LD information is not available, which is a particular concern in broad-ancestry studies or meta analyses, or when speed is particularly important, that RápidoPGS-single is chosen. Our method for improving the speed of the SuSiE, by filtering SNPs according to p value, is approximate and was not recommended by the method authors. We use it to allow this fine mapping approach to run in a reasonable time for our purposes, and it appears to perform well in this situation, but it is unlikely to be optimal. It is possible a more accurate PGS could be constructed if all SNPs were supplied, but this would be at the cost of more than an order of magnitude slower speed.

Like all methods used here for comparison, RápidoPGS-multi requires LD matrices, constructed on the same or similar population to the dataset on which the PGS is trained. However, since individual data is often not available due to privacy concerns, this can be done using a publicly available reference panel. Despite its relatively small size (2504 individuals of worldwide origin), 1000 Genomes Project Phase III is publicly available, and as we have shown, can be used for LD matrix computation and obtain good results. We provide a function to download and pre-process a 1000 Genomes-based reference panel from scratch, although users are free to use their own panel.

We are not the first to suggest that fine mapping approaches can be helpful for PGS construction, Newcombe and colleagues (2019) used reversible jump MCMC to fit a fine mapping model to GWAS summary statistics, parallelising across LD blocks as we do here. However, rather than estimating posterior inclusion probabilities and using these to shrink frequentist effect estimates, they averaged samples from the posterior distribution of causal effect estimates in their fine mapping model, thus mirroring the PGS approach of focusing on the true effect estimates. Our work offers an alternative method to generate the weights *w_i_*. Other advances in fine mapping methods may be transferable to the PGS setting. For example, PGS are known to have less predictive power in populations other than that used for training (Martin *et al.*, 2019). This is a particular issue given the eurocentric focus of GWAS to date. Restricting PGS to SNPs with known functional annotations has been shown to increase the portability of scores between ancestries (Amariuta *et al.*, 2020), but relevant functional annotations can be incomplete. Fine mapping offers a natural means to incorporate levels of annotation data through variable per-SNP priors, and methods have been developed to learn appropriate priors through hierarchical Bayesian approaches (Pickrell, 2014). Alternatively, established trans-ancestry fine mapping approaches may be useful (Morris, 2011). Thus, while our work presents a method designed for fast and easy generation of PGS, it also highlights that the current challenges for PGS may be potentially addressed through adaptation of fine-mapping approaches which addressed similar challenges in that field.

## Acknowledgements

We thank Florian Privé and Bjarni Vilhjálmsson for useful guidance on running LDpred2. We thank Chris Q. Eijsbouts for valuable feedback on early versions of the RápidoPGS package.

## Funding information

This research was funded by the by the MRC (MC_UU_00002/4) and the Wellcome Trust [WT107881, 203950/Z/16/A], and supported by the NIHR Cambridge BRC (BRC-1215-20014). The views expressed are those of the author(s) and not necessarily those of the NHS, the NIHR or the Department of Health and Social Care. This research was funded in whole, or in part, by the Wellcome Trust [WT107881, 203950/Z/16/A]. For the purpose of open access, the author has applied a CC BY public copyright licence to any Author Accepted Manuscript version arising from this submission.

This research has been conducted using the UK Biobank Resource under Application Number 30931.

## Web resources

R - https://cran.r-project.org/

RápidoPGS - https://github.com/GRealesM/RapidoPGS

LDpred2 - https://privefl.github.io/bigsnpr/articles/LDpred2.html

LD detect datasets - https://bitbucket.org/nygcresearch/ldetect-data/src

GWAS catalog - https://www.ebi.ac.uk/gwas/

1000 Genomes Project - https://www.internationalgenome.org/

LDHub - http://ldsc.broadinstitute.org/ldhub/

## Data and code availability

Summary statistics datasets used are publicly available in their respective publications (see Suppl. Table 1). Code used in the analyses is available at GitHub (https://github.com/GRealesM/RapidoPGS_paper) and code used for model evaluation is available at GitLab (https://gitlab.com/evigorito/applyrapidopgs).

We have extensively used tools in the *bigsnpr* (Privé *et al.*, 2018) and *data.table* for large dataset handling and analysis.

## Description of Supplemental Data

Supplemental data contains a note, 10 figures, and 5 tables.

**Supplementary Figure 1.**
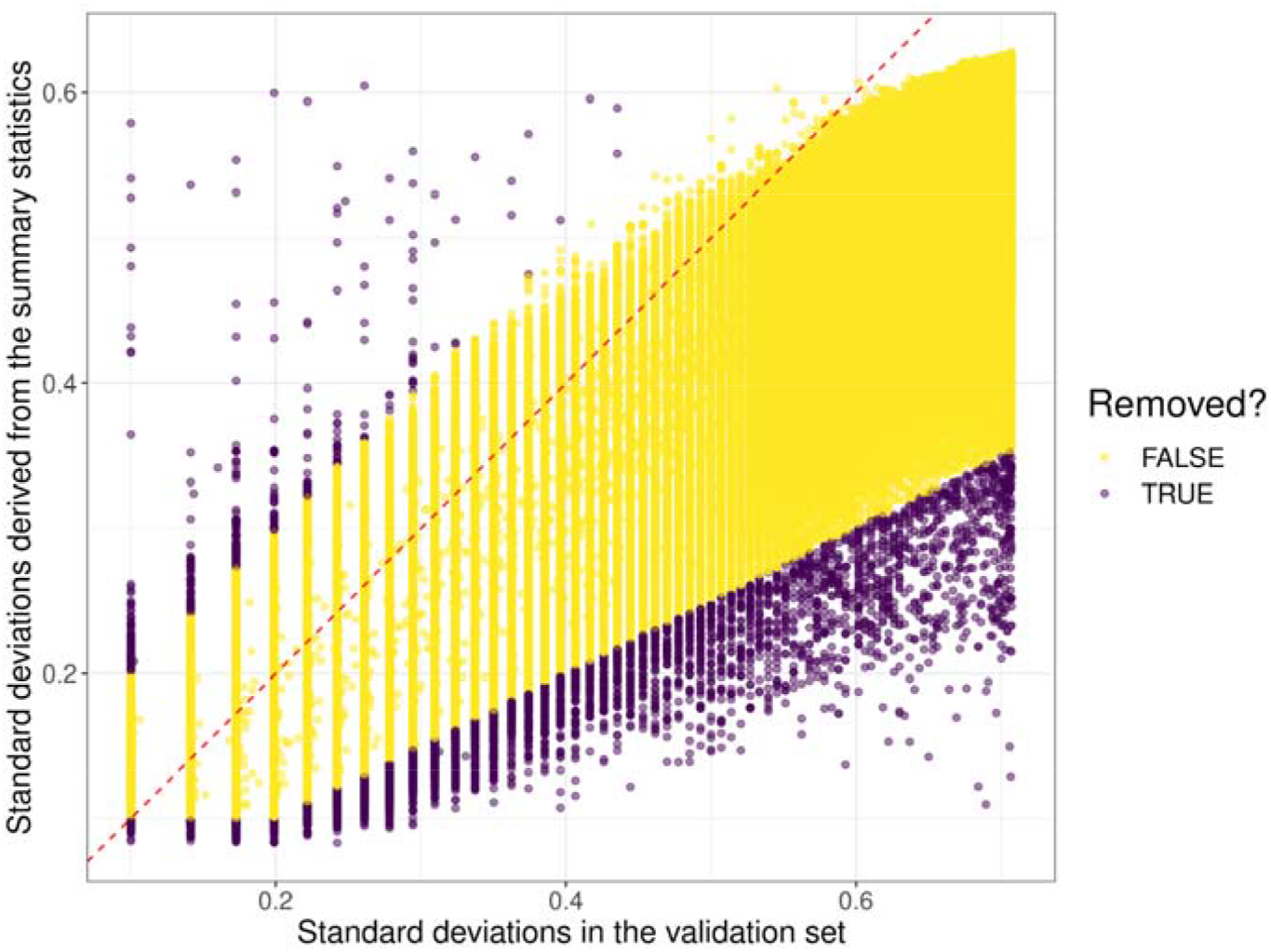
QC plot for Asthma, showing included and excluded SNPs by their SDss (y-axis) and SDval (x-axis) values. The dotted line corresponds to the x = y line.

**Supplementary Figure 2.**
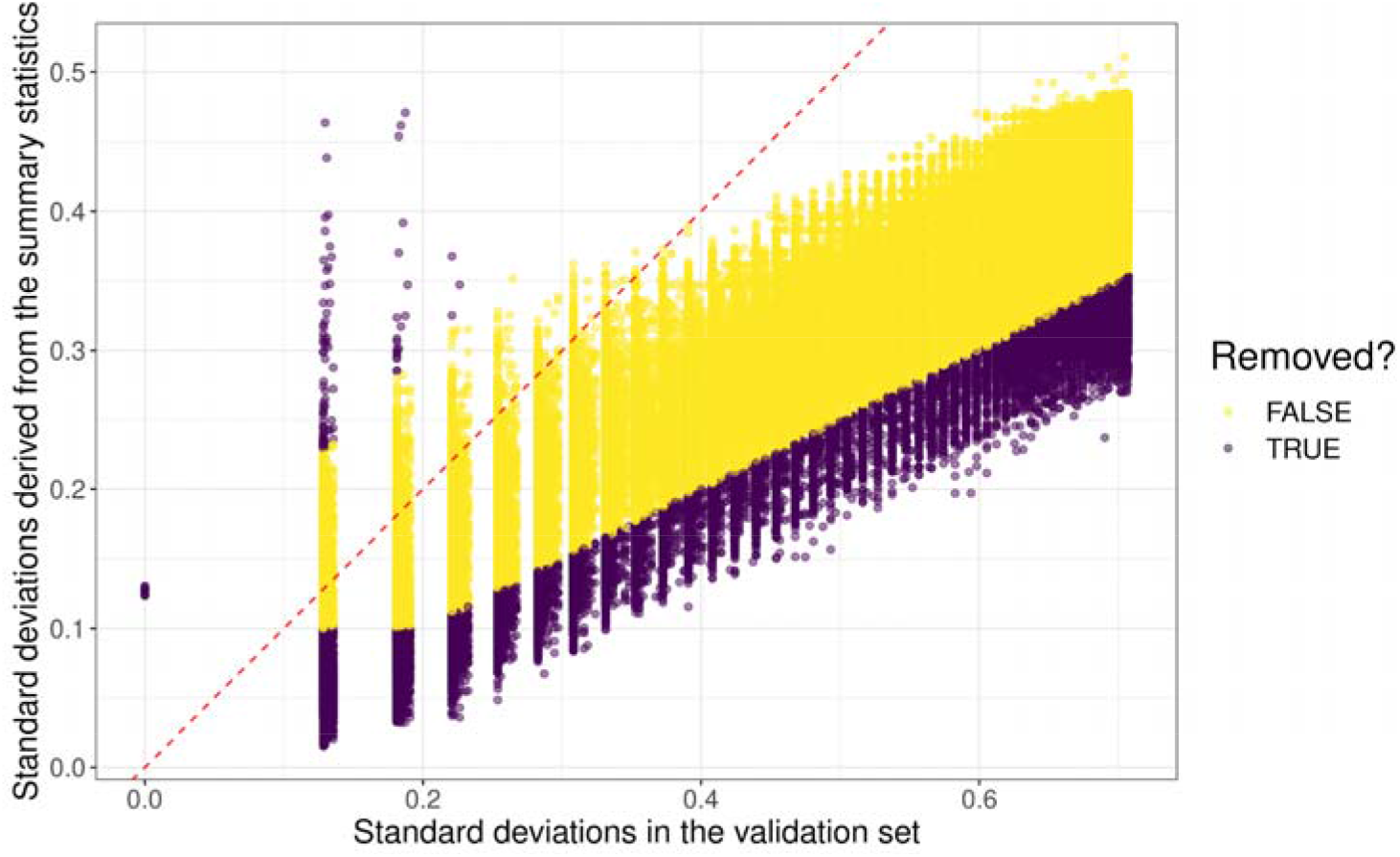
QC plot for BMI, showing included and excluded SNPs by their SDss and SDval values. The dotted line corresponds to the x = y line.

**Supplementary Figure 3.**
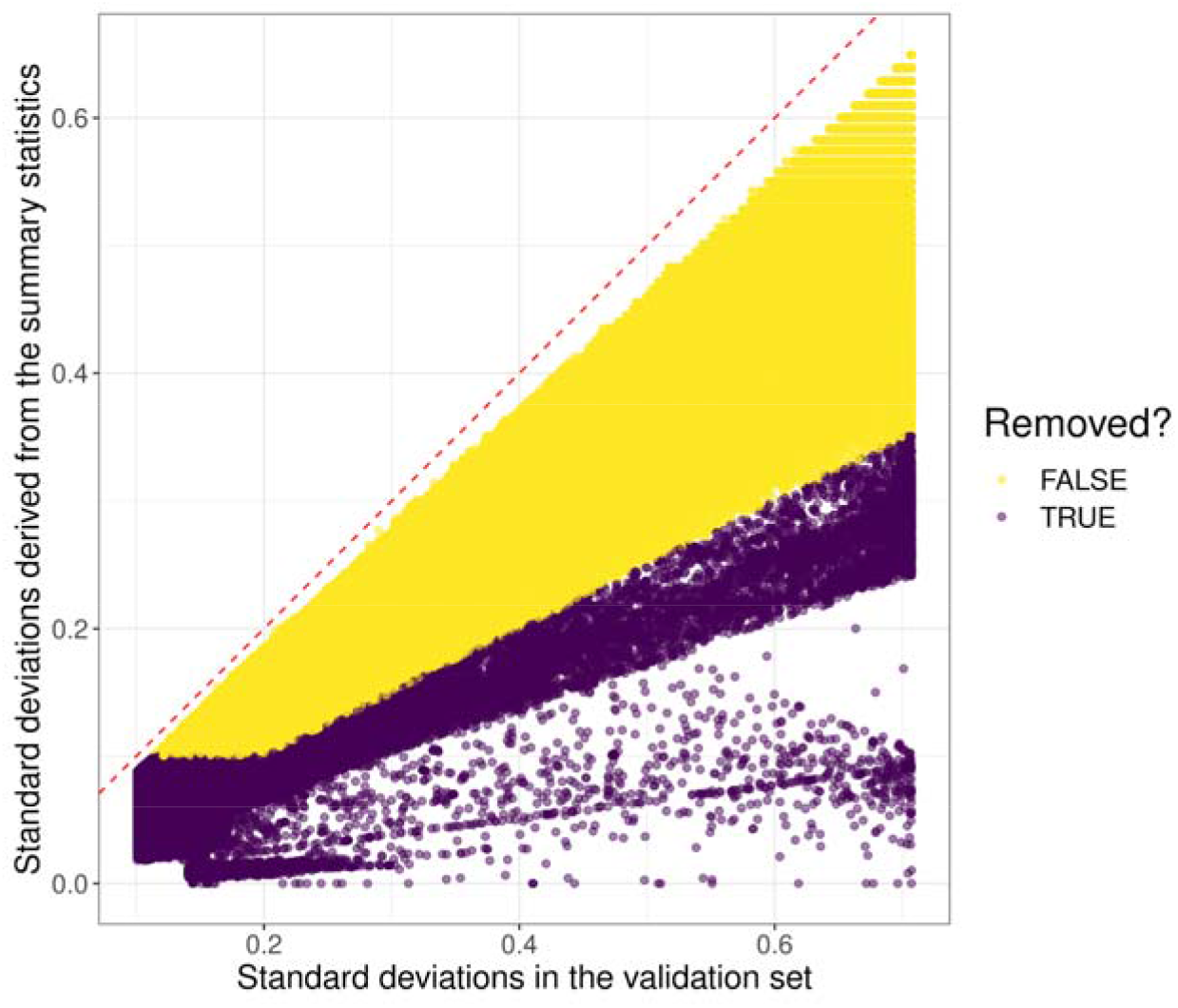
QC plot for BRCA, showing included and excluded SNPs by their SDss and SDval values.The dotted line corresponds to the x = y line.

**Supplementary Figure 4.**
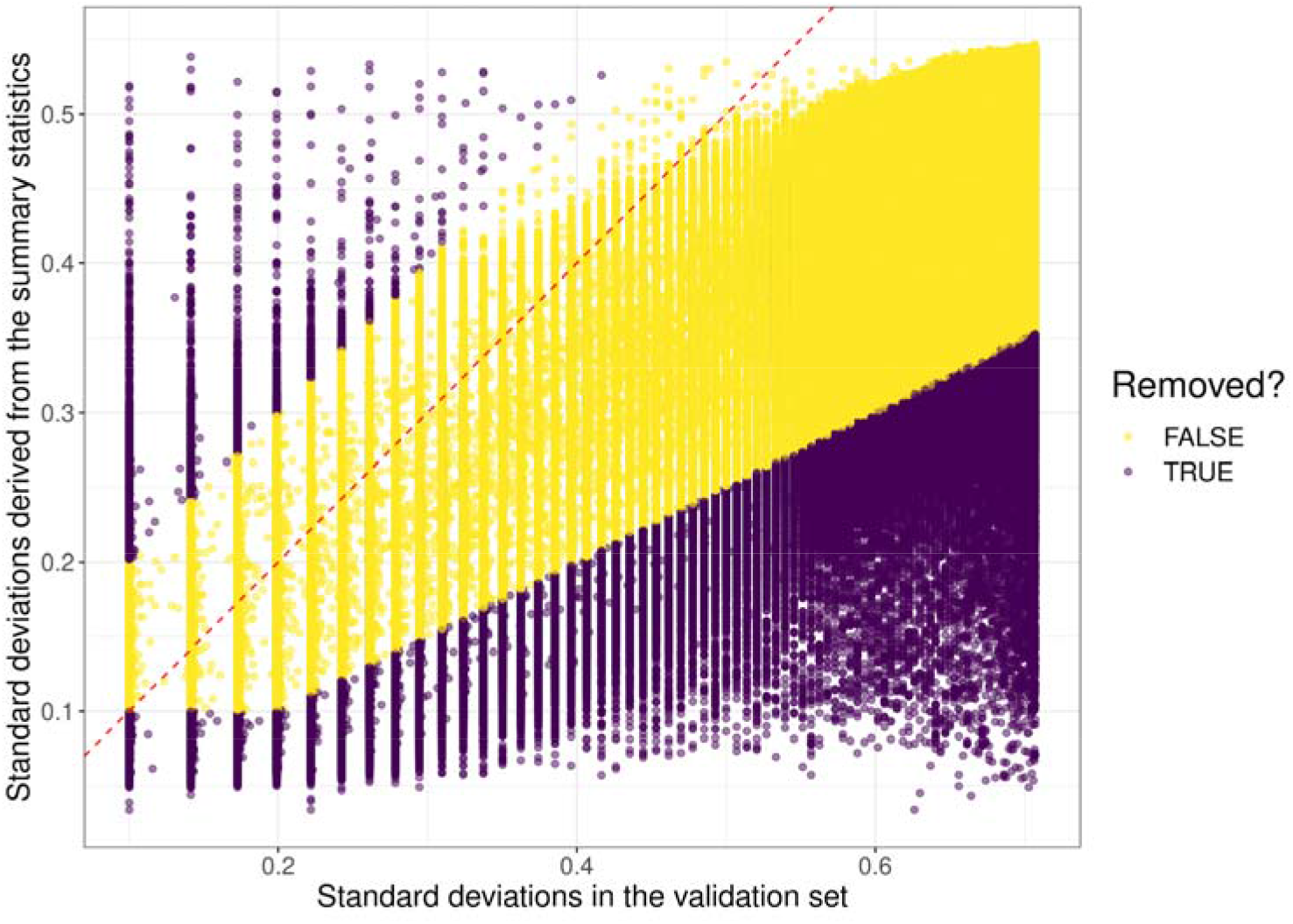
QC plot for CAD, showing included and excluded SNPs by their SDss and SDval values. The dotted line corresponds to the x = y line.

**Supplementary Figure 5.**
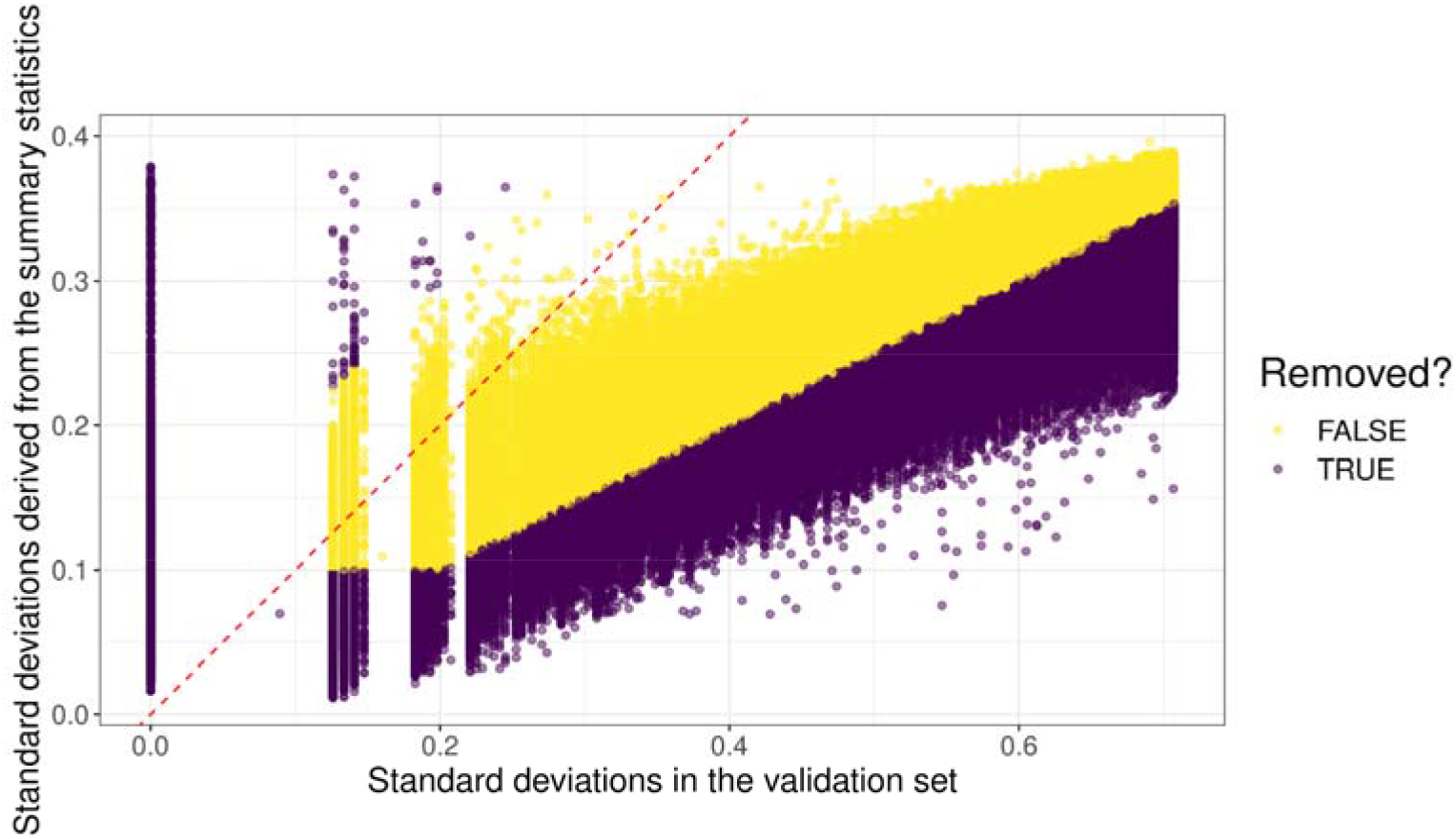
QC plot for Height, showing included and excluded SNPs by their SDss and SDval values. The dotted line corresponds to the x = y line.

**Supplementary Figure 6.**
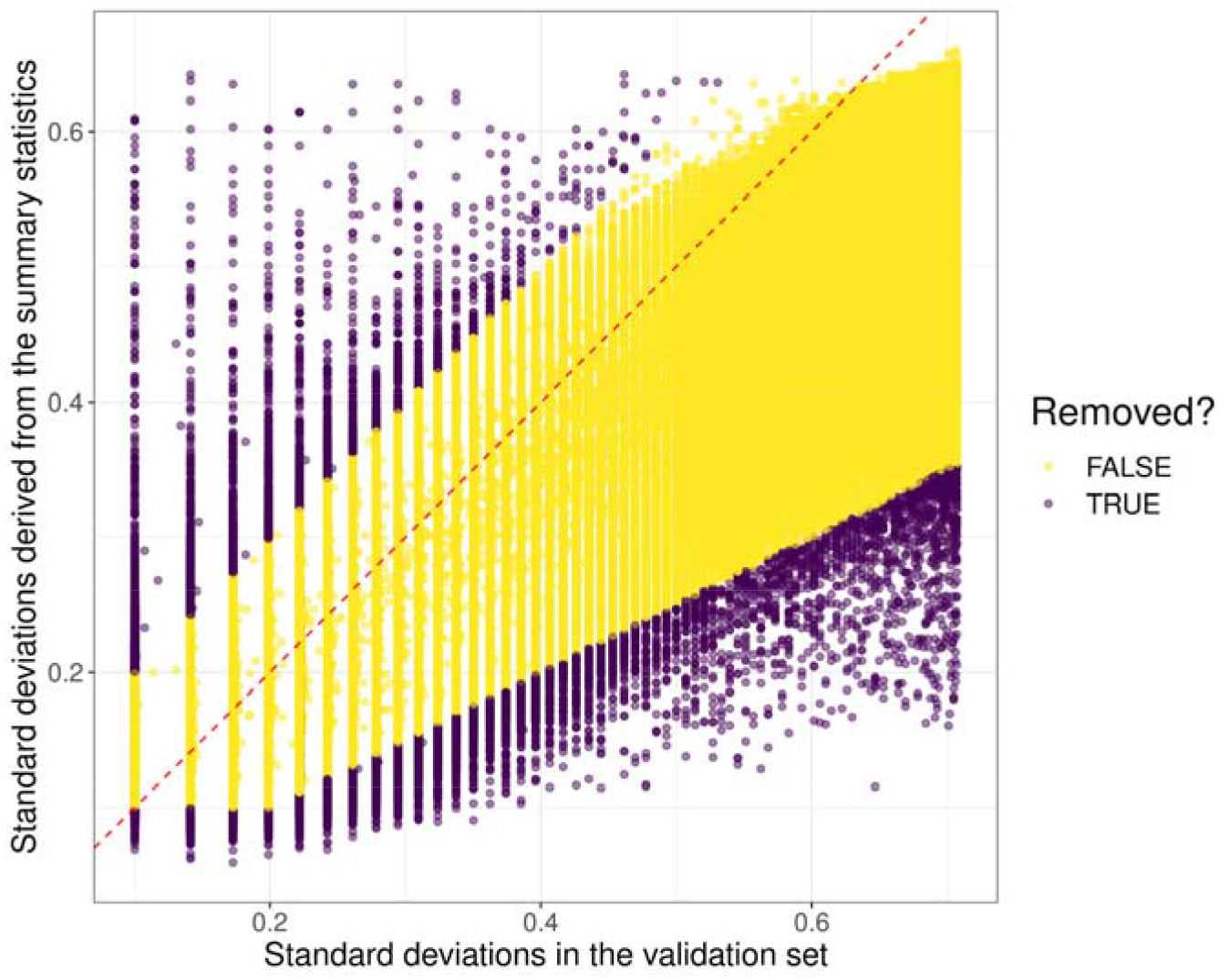
QC plot for MDD, showing included and excluded SNPs by their SDss and SDval values. The dotted line corresponds to the x = y line.

**Supplementary Figure 7.**
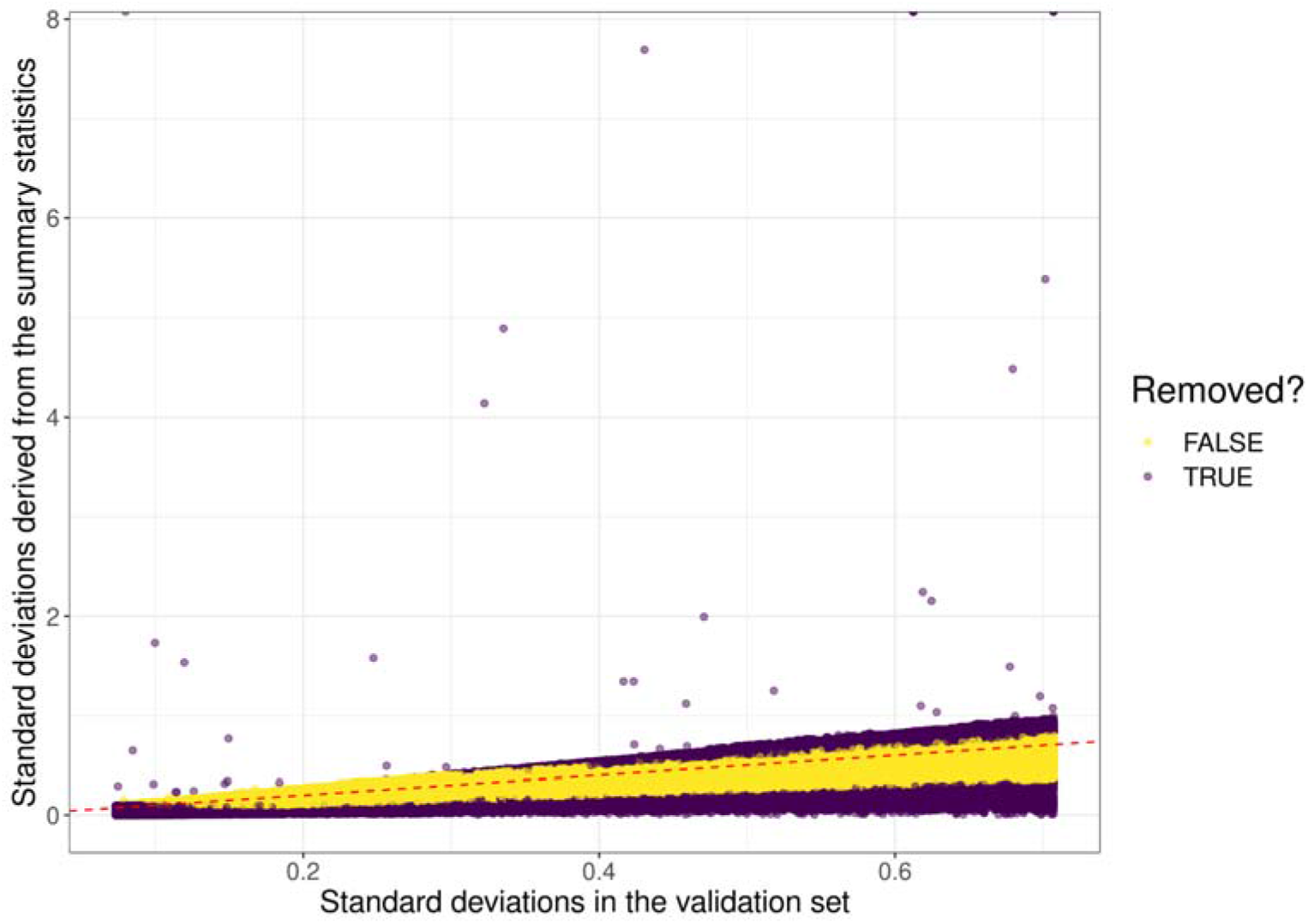
QC plot for PRCA, showing included and excluded SNPs by their SDss and SDval values. The dotted line corresponds to the x = y line.

**Supplementary Figure 8.**
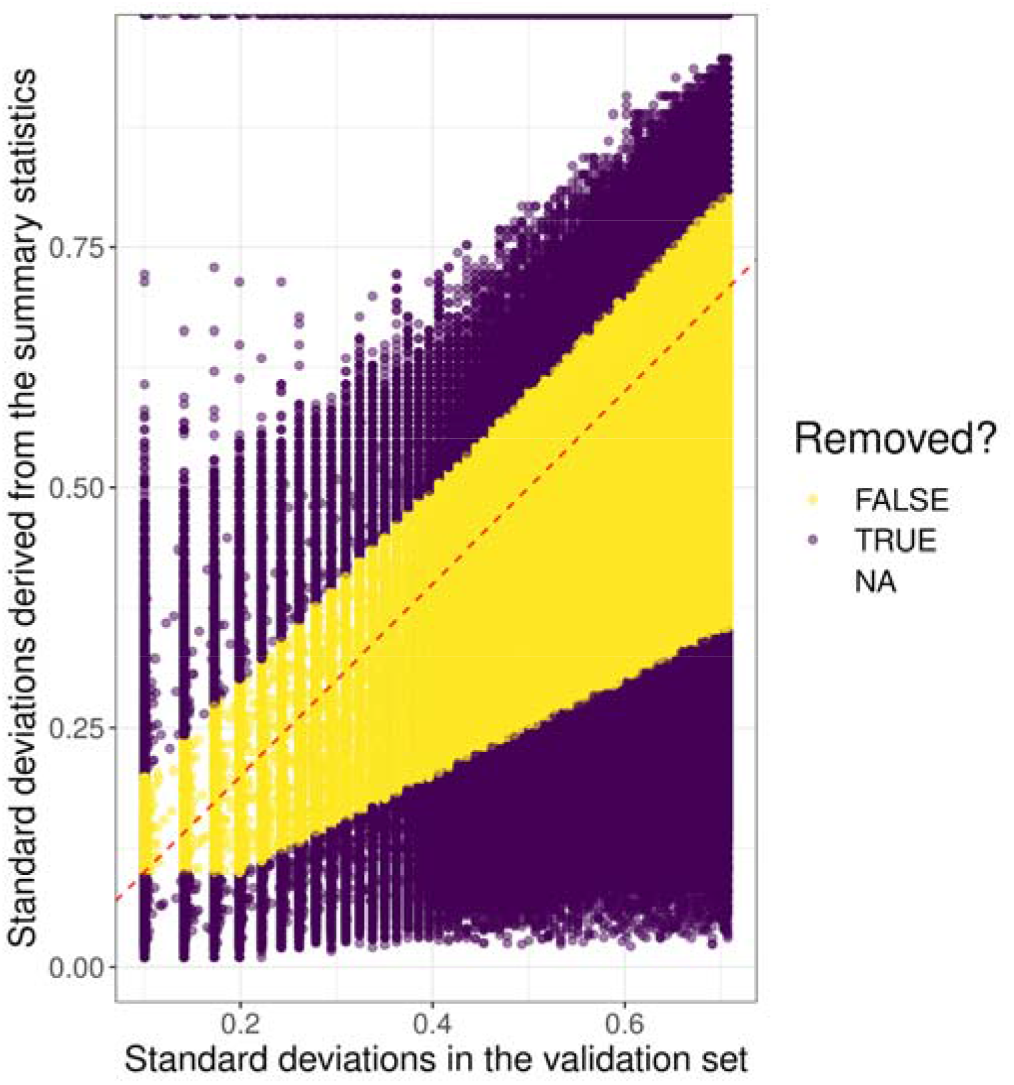
QC plot for RA, showing included and excluded SNPs by their SDss and SDval values. The dotted line corresponds to the x = y line.

**Supplementary Figure 9.**
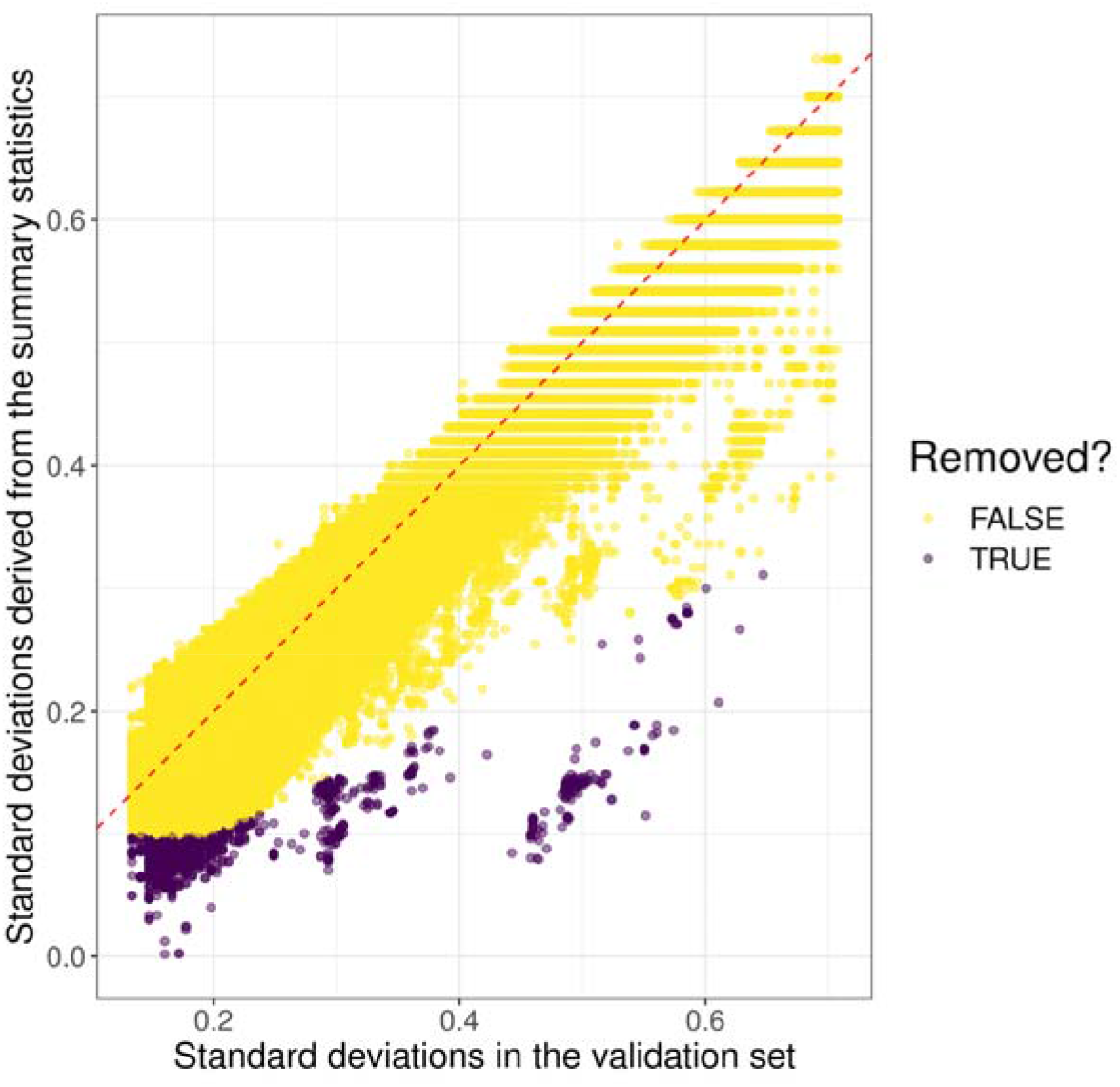
QC plot for T1D, showing included and excluded SNPs by their SDss and SDval values.The dotted line corresponds to the x = y line.

**Supplementary Figure 10.**
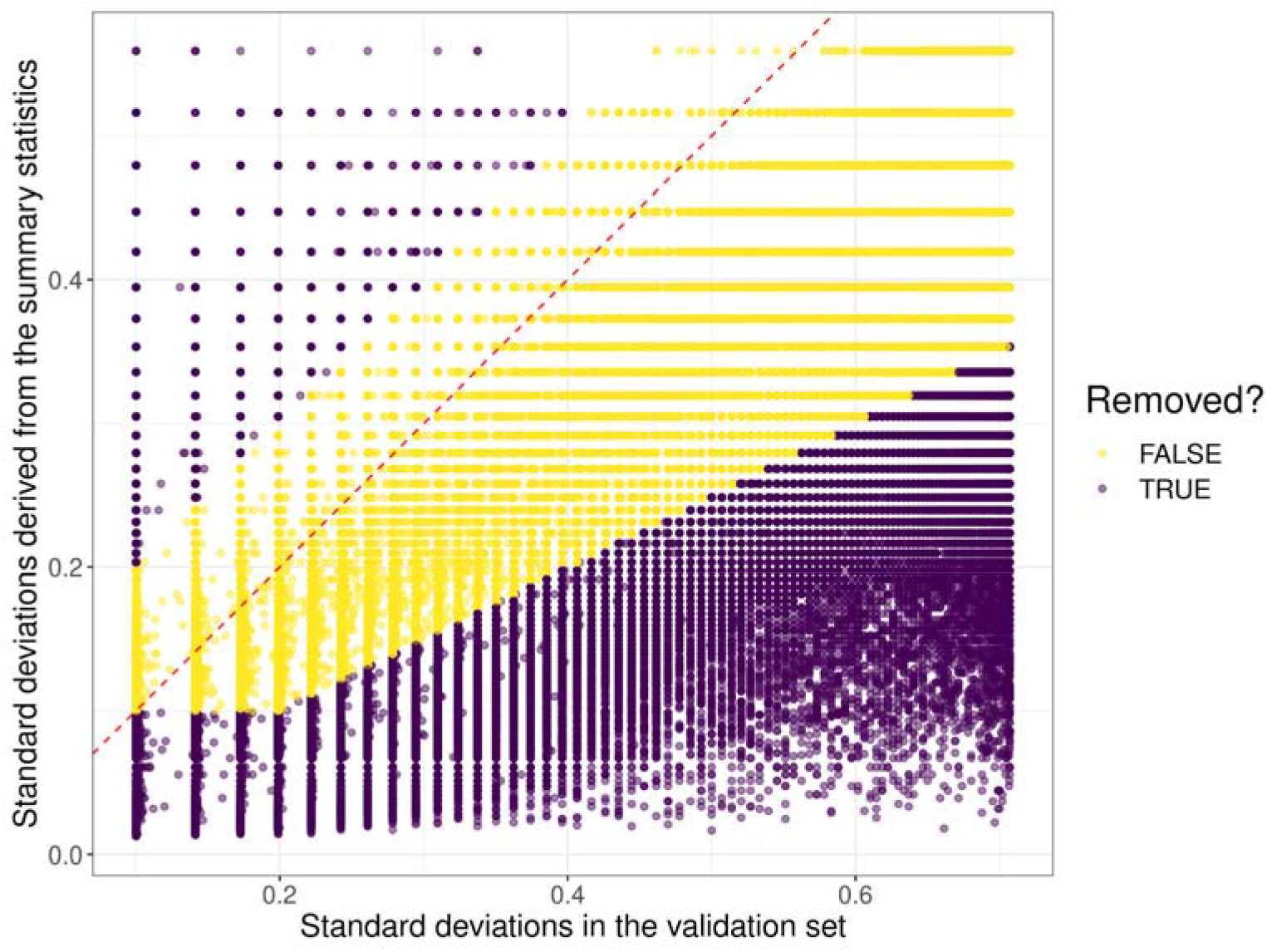
QC plot for T2D, showing included and excluded SNPs by their SDss and SDval values. The dotted line corresponds to the x = y line.

## Supplementary Table legends

**Supplementary Table 1.** GWAS summary statistic datasets. For T1D dataset, we used Cooper et al., 2017, instead of Censin et al., 2017 dataset. These should be equivalent so we provide a link to the available Censin et al. dataset.

Supplementary Table . Results of PGS model evaluations model evaluations, in AUC and r^2^.

Supplementary Table 3. Results for PGS benchmark for all tests, in seconds.

Supplementary Table 4. r^2^ differences between RápidoPGS approaches, LDpred2, PRScs, and SBayesR, using HapMap3 variants. Positive values mean RápidoPGS performed better (ie. achieved larger r^2^) than the compared method.

Supplementary Table 5. r^2^ ratio between RápidoPGS approaches, LDpred2, PRScs, and SBayesR, using HapMap3 variants. Values larger than one mean RápidoPGS performed better (ie. achieved larger r^2^) than the compared method.

