## Supplemental Note for "RápidoPGS: A rapid polygenic score calculator for summary GWAS data without a test dataset"

Assume we have a trait

$$Y = \sum_{i=1}^{m} X_i \beta_i + \epsilon$$

with $\beta_i \sim N(0, W)$ and the sum taken over $m$ SNPs that are causal for the trait. Heritability is

$$h^2 = \frac{\operatorname{Var}(Y - \epsilon)}{\operatorname{Var} Y} \Rightarrow \operatorname{Var} \epsilon = (1 - h^2) \operatorname{Var}(Y)$$

Then it follows that

$$\begin{aligned} \operatorname{Var}(Y) &= \operatorname{Var}(\sum_{i=1}^{m} X_i \beta_i) + \operatorname{Var}(\epsilon) \\ &= \sum_{i=1}^{m} \beta_i^2 \operatorname{Var}(X_i) + (1 - h^2) \operatorname{Var}(Y) \end{aligned}$$

Here Var is taken across individuals, and $\beta_i$ is fixed but unknown for each SNP $i$. We replace $\beta_i^2$ by its expected value across SNPs, W. $\sum \operatorname{Var}(X_i)$ is also unknown, but we have estimates of $\operatorname{Var}(\hat{\beta}_i) = \frac{\operatorname{Var} Y}{N_i \operatorname{Var} X_i}$ where $N_i$ is the number of individuals with data at SNP $i$. Therefore

$$\sum_{i=1}^{m} \operatorname{Var}(X_i) \simeq \sum_{i=1}^{m} \frac{\operatorname{Var} Y}{N_i \operatorname{Var}(\hat{\beta}_i)} = \operatorname{Var} Y \sum_{i=1}^{m} \frac{1}{N_i \operatorname{Var}(\hat{\beta}_i)}$$

where the sum is taken across all $m$ causal SNPs. We do not know what the causal SNPs are, but assume instead that $N_i, \operatorname{Var} \hat{\beta}_i$ take the same distribution across causal SNPs $1, \ldots, m$ and non-causal SNPs $m + 1, \ldots, p$. Then we can estimate

$$\sum_{i=1}^{m} \frac{1}{N_i \operatorname{Var} \hat{\beta}_i} \simeq \frac{m}{p} \sum_{i=1}^{p} \frac{1}{N_i \operatorname{Var} \hat{\beta}_i}$$

Finally, $m$ is still unknown, but we have already specified the fraction of truly causal SNPs we expect as $\pi$ so we estimate $\widehat{(\frac{m}{p})} = \pi$. Thus, putting it all together, we can write

$$\begin{aligned} \operatorname{Var}(Y) &= W \sum_{i=1}^{m} \operatorname{Var}(X_i) + (1 - h^2) \operatorname{Var}(Y) \\ &\simeq W \operatorname{Var}(Y) \pi \sum_{i=1}^{p} \frac{1}{N_i \operatorname{Var} \hat{\beta}_i} + (1 - h^2) \operatorname{Var}(Y) \end{aligned}$$

so that

$$W \simeq h^2 \Big/ \pi \sum_{i=1}^{p} \frac{1}{N_i \operatorname{Var} \beta_i}$$
